# Discovery of a novel small protein factor involved in the coordinated degradation of phycobilisomes in cyanobacteria

**DOI:** 10.1101/2020.06.14.150680

**Authors:** Vanessa Krauspe, Matthias Fahrner, Philipp Spät, Claudia Steglich, Nicole Frankenberg-Dinkel, Boris Macek, Oliver Schilling, Wolfgang R. Hess

## Abstract

Phycobilisomes are the major pigment-protein antenna complexes that perform photosynthetic light harvesting in cyanobacteria, rhodophyte and glaucophyte algae. Up to 50% of the cellular nitrogen can be stored in their giant structures. Accordingly, upon nitrogen depletion, phycobilisomes are rapidly degraded. This degradation is tightly coordinated, follows a genetic program and involves small proteins serving as proteolysis adaptors. Here, we describe the role of NblD, a novel factor in this process in cyanobacteria. NblD is a cysteine-rich, 66-amino acid small protein that becomes rapidly induced upon nitrogen starvation. Deletion of the *nblD* gene in the cyanobacterium *Synechocystis* prevents the degradation of phycobilisomes, leading to a nonbleaching (*nbl*) phenotype. Competition experiments provided direct evidence for the physiological importance of NblD. Complementation by a plasmid-localized gene copy fully restored the phenotype of the wild type. Overexpression of NblD under nitrogen-replete conditions showed no effect, in contrast to the unrelated proteolysis adaptors NblA1 and NblA2, which can trigger phycobilisome degradation ectopically. Transcriptome analysis revealed that nitrogen starvation correctly induces *nblA1/2* transcription in the Δ*nblD* strain implying that NblD does not act as a transcriptional (co-)regulator. However, fractionation and coimmunoprecipitation experiments indicated the presence of NblD in the phycobilisome fraction and identified the β-phycocyanin subunit as its target. These data add NblD as a new factor to the genetically programmed response to nitrogen starvation and demonstrate that it plays a crucial role in the coordinated dismantling of phycobilisomes when nitrogen becomes limiting.

**Significance Statement:** During genome analysis, genes encoding small proteins are frequently neglected. Accordingly, small proteins have remained underinvestigated in all domains of life. Based on a previous systematic search for such genes, we present the functional analysis of the small protein NblD in a photosynthetic cyanobacterium. We show that NblD plays a crucial role during the coordinated dismantling of phycobilisome light-harvesting complexes. This disassembly is triggered when the cells run low in nitrogen, a condition that frequently occurs in nature. Similar to the NblA proteins that label phycobiliproteins for proteolysis, NblD binds to phycocyanin polypeptides but has a different function. The results show that, even in a well-investigated process, crucial new players can be discovered if small proteins are taken into consideration.

## Introduction

### The response of cyanobacteria to nitrogen starvation is governed by a complex genetic program

Nitrogen is an essential element of all organisms and frequently the main nutrient limiting life of photoautotrophic primary producers in many terrestrial, freshwater and marine ecosystems (1, 2). Cyanobacteria are the only prokaryotic primary producers performing oxygenic photosynthesis. Some cyanobacteria are of overwhelming relevance for the global biogeochemical cycles of carbon and nitrogen, exemplified by the marine *Prochlorococcus* and *Synechococcus* with their estimated global mean annual abundances of 2.9 ± 0.1 ×10^27^ and 7.0 ± 0.3 × 10^26^ cells (3–5). Other cyanobacteria came into focus for the ease of their genetic manipulation, fast growth and as platforms for the CO_2_-neutral production of diverse valuable products (e.g., (6, 7)). With regard to their response to nitrogen starvation, cyanobacteria can be divided into two major physiological groups. Diazotrophic genera such as *Trichodesmium, Nodularia, Cyanothece, Nostoc* or *Anabaena* avoid nitrogen limitation by expressing nitrogenase to fix the omnipresent gaseous N_2_. In contrast, non-diazotrophic cyanobacteria such as *Synechococcus* and *Synechocystis* stop growth and switch their metabolism from anabolism to maintenance, a process that is controlled by a complex genetic program (8–10). An acute scarcity in the available nitrogen is sensed directly by two central regulators of nitrogen assimilation in cyanobacteria, P_II_ and NtcA, by binding the key metabolite 2-oxoglutarate (2-OG) (11–15). 2-OG is the substrate for amination by glutamine oxoglutarate aminotransferase (GOGAT), which catalyzes the transfer of the amino group from glutamine to 2-OG, yielding two molecules of glutamate. This glutamate then is the substrate for amination by glutamine synthetase, the central enzyme of nitrogen assimilation in cyanobacteria. As a consequence, the intracellular level of 2-OG starts to increase once these reactions slow down because nitrogen is becoming scarce, making 2-OG an excellent indicator of nitrogen status (16). Upon binding 2-OG, the activity of NtcA, the main transcriptional regulator of nitrogen assimilation, becomes stimulated (11, 12). Depending on the 2-OG level, PipX switches from binding to P_II_ to interacting with NtcA enhancing the binding affinity of this complex to target promoters further (13, 15). In the model cyanobacterium *Synechocystis* sp. PCC 6803 (from here on *Synechocystis* 6803), NtcA directly activates 51 genes and represses 28 other genes after 4 h of nitrogen starvation (17). Among the NtcA-activated genes are the cotranscribed genes *nblA1* and *nblA2* (*ssl0452* and *ssl0453*) (17). Once they are expressed, NblA proteins impact the physiology dramatically. They initiate the degradation of phycobiliproteins, the major light harvesting pigments in the cyanobacterial phycobilisomes, a process that is visible by the naked eye because it leads to a color change from blue-green to yellowish.

### Phycobilisomes, the most efficient structures for photosynthetic light harvesting, become degraded during nitrogen starvation

Phycobilisomes are the major light-harvesting system in red algae and most cyanobacteria. Phycobilisomes are giant protein-pigment complexes anchored to the thylakoid membranes (18, 19) that absorb light mainly in the “green gap” between 580 and 650 nm. Phycobilisome complexes are highly abundant and may contain up to 50% of the soluble cellular protein and nitrogen content (20). Therefore, their ordered disassembly is part of the physiological response to nitrogen starvation (10, 21–24) that can free a substantial amount of amino acids and release the nitrogen bound as part of the photosynthetic pigment molecules.

The degradation of phycobiliproteins is initiated by NblA proteins. Mutations in the *nblA* (*nonbleaching* A) genes yield, unlike the wild-type strain, a nonbleaching phenotype under nitrogen starvation because they do not degrade their phycobilisomes (25–27). Binding experiments indicated that NblA likely interacts with the α-subunits of phycobiliproteins in *Tolypothrix* PCC 7601 (28) and *Anabaena* sp. PCC 7120 (29); however, in *Synechococcus elongatus* UTEX 2973, it was found to bind to the N-terminus of β-phycocyanin (30). Pull-down experiments then led to the discovery that NblA acts as an adaptor protein for the Clp protease by also interacting with the ClpC chaperone (31). Because the chaperone partner determines the substrate specificity of this protease, NblA presents the protein components of the phycobilisome for proteolysis. The extensive characterization of additional mutants with an *nbl* phenotype in *Synechococcus elongatus* PCC 7942 (*S. elongatus* 7942) led to the discovery of further enzymes and regulatory proteins, NblB1, NblB2, NblC, NblR and NblS, which play roles in the preprogrammed disassembly of phycobilisomes during nitrogen starvation (**Table 1**).

**Table 1.** Proteins previously identified in different cyanobacteria as involved in the programmed phycobilisome disassembly and their homologs in *Synechocystis* 6803. Synonymous names of certain proteins are separated by a slash.

| Protein name | Gene ID | Function | Reference |
| --- | --- | --- | --- |
| NblA1 | ssl0452 | protease adaptor | (32) |
| NblA2 | ssl0453 | protease adaptor | (32) |
| ClpC | sll0020 | HSP100 chaperone partner of Clp protease | (70) |
| NblB1 | sll1663 | bilin lyase homolog | (32, 63) |
| NblB2 | slr1687 | bilin lyase homolog | (32, 63) |
| NblC | sll1968 | regulator | (88) |
| NblR | sll0396 | regulator | (26) |
| GntR | sll1961 | regulator | (69) |
| Hik33/<br>NblS/DspA | sll0698 | sensor kinase | (33, 89) |
| RpaB/Ycf27 | slr0947 | response regulator | (90) |

In contrast to these observations in *S. elongatus* 7942, the mutations of corresponding genes in *Synechocystis* 6803, Δ*slr1687* (NblB homolog #2), Δ*sll0396* (NblR homolog) and Δ*sll0698* (NblS homolog), did not yield a nonbleaching phenotype during nitrogen depletion (32, 33). These findings suggest that, in addition to commonalities, substantial differences exist in the response of certain species to nitrogen depletion and the organization of the photosynthetic apparatus. Nevertheless, the program governing the acclimation of non-diazotrophic cyanobacteria to nitrogen starvation and the process leading to the ordered degradation of phycobilisomes and the photosynthetic pigments therein is considered as well understood.

### Small proteins in Cyanobacteria

The afore mentioned proteolysis adaptors NblA1 and NblA2 are small proteins of just 62 and 60 residues, respectively. During standard genome analyses, small open reading frames (smORFs) encoding such proteins shorter than 70 amino acids are frequently neglected. The identification of small proteins by mass spectrometry (MS) based on shotgun proteomics is also difficult. According to their length, these proteins contain only a few or even miss cleavage sites for commonly used proteases, such as trypsin. Consequently, the number of generated peptides is smaller than that for larger proteins. Additionally, MS spectra of low abundant proteins with only a few unique peptides might not fulfill common quality criteria and are removed during filtering. Thus, the number of genes encoding small proteins has been systematically underestimated. In strong contrast is the finding that genes encoding small proteins constitute an essential genomic component in bacteria (34). Recent ribosome profiling studies in bacteria using the inhibitor of translation retapamulin suggest that a high number of previously unexplored small proteins exist in bacteria (35, 36). Detailed analyses are required to identify their functions at the molecular level.

Cyanobacteria provide a paradigm for small protein functions also in addition to the known NblA functions. Extensive work on the photosynthetic apparatus lead to the functional characterization of 19 small proteins with even fewer than 50 amino acids. These play indispensable roles in photosystem II (genes *psbM, psbT* (*ycf8*), *psbI, psbL, psbJ, psbY, psbX, psb30* (*ycf12*), *psbN, psbF, psbK* (37, 38)), photosystem I (*psaM, psaJ, psaI* (39)), photosynthetic electron transport (Cyt*b*_6_*f* complex; *petL, petN, petM, petG* (40–42)), and photosynthetic complex I (*ndhP, ndhQ* (43)) or have accessory functions (*hliC* (*scpB*) (44)). The shortest annotated photosynthetic protein conserved in cyanobacteria has 29 amino acids — the cytochrome *b*_6_*f* complex subunit VIII, encoded by *petN* (45).

We have previously analyzed the primary transcriptomes of the model cyanobacteria *Synechocystis* sp. PCC 6803 (from here *Synechocystis* 6803) and the closely related strain *Synechocystis* sp. PCC 6714 (46–48). Based on these analyses, several smORFs likely encoding previously unknown small proteins were computationally predicted. Experiments using a small 3xFLAG epitope tag fused in-frame to the second-to-last codon of the smORF under control of their native promoters and 5’UTRs validated five of these smORFs to encode small proteins (49).

Here, we analyzed one of these small proteins, the 66-residues NsiR6 (nitrogen stress-induced RNA 6), which is highly upregulated following nitrogen removal (46, 49). We show that this small protein is a crucial factor in the genetically programmed response to nitrogen starvation executing a previously unrecognized role in the coordinated disassembly of phycobilisomes. Based on the observed *nbl* phenotype, we renamed NsiR6 to NblD.

## Results

### Homologs of *nblD* are widely conserved within the cyanobacterial phylum

The *nblD* gene in *Synechocystis* 6803 is located on the chromosome, between the genes *slr1704* encoding a hypothetical S-layer protein and *sll1575* encoding the serine-threonine protein kinase SpkA (50), at a distance of less than 2.5 kb from the *cpcBAC2C1D* operon encoding phycobilisome proteins (**Fig. 1A**). Database searches identified 176 homologs of *nblD* in species belonging to all morphological subsections except section V (*Fischerella* and other cyanobacteria featuring filaments with a branching morphology). Homologous genes were also detected in the three available chromatophore genomes of photosynthetic *Paulinella* species (52). The endosymbiont chromatophore genomes have been reduced to about one-third the size of the genome of its closest free-living relatives (53). Hence, the presence of *nblD* homologs in chromatophore genomes indicates a possible important function connected to the remaining sections of its metabolism. We detected putative homologs also in two diatom-associated symbionts, *Calothrix rhizosoleniae* and *Richelia intracellularis*. However, we found no homologs in the genomes of UCYN-A “Candidatus Atelocyanobacterium thalassa” endosymbionts, which have the capacity for nitrogen fixation but lack photosystem II and phycobilisomes (54, 55), and in the well-studied model *S. elongatus* 7942. Homologs of NblD are also lacking in the genera *Prochlorococcus* and most *Acaryochloris*, which use alternative light-harvesting mechanisms. We noticed, however, the presence of a homolog in *Acaryochloris thomasi* RCC1774, an isolate with a very different pigmentation (56).

**Fig. 1.**
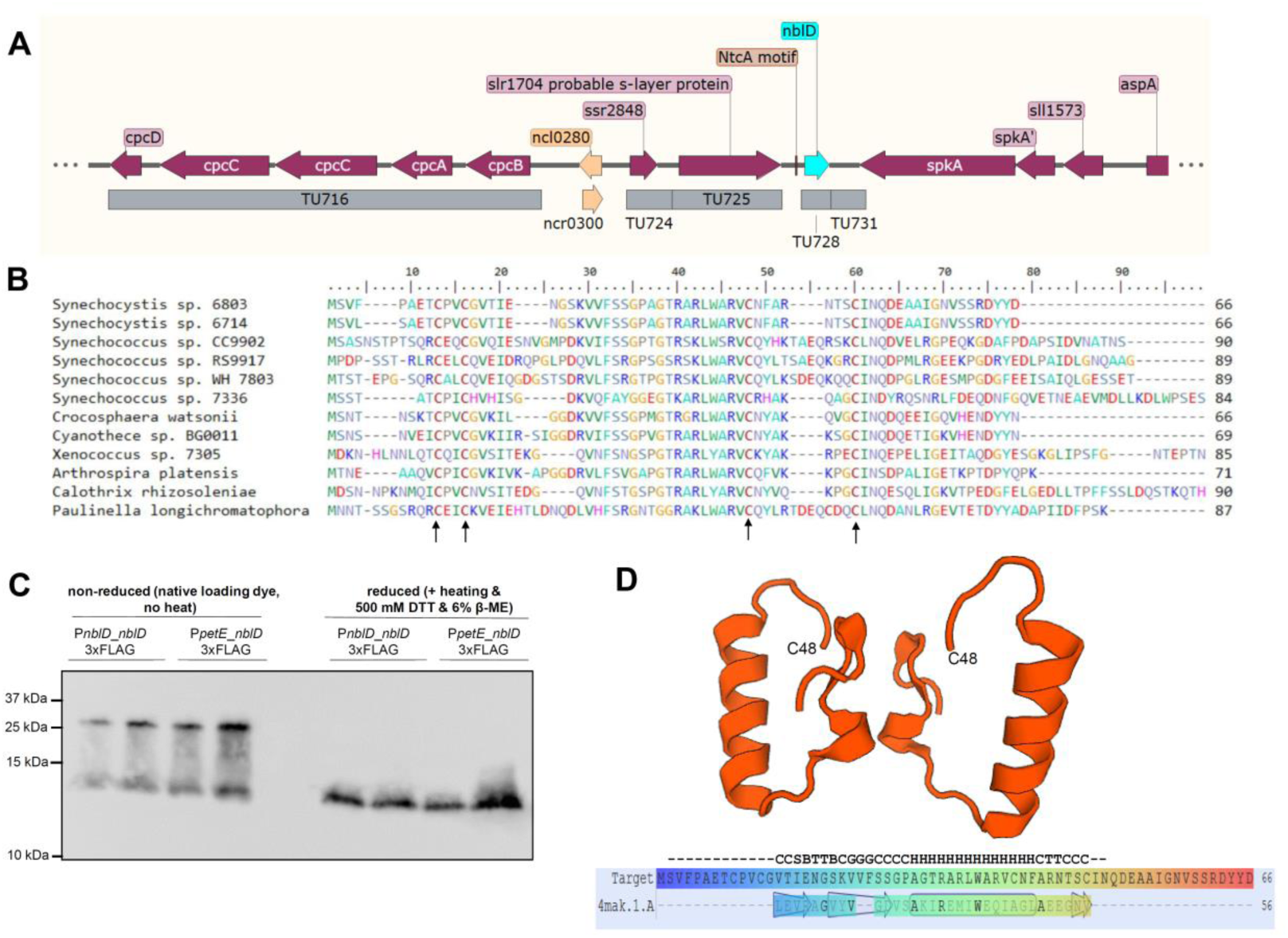
**A**. Genomic location of *nblD* in *Synechocystis* 6803. Transcriptional units (TUs) are indicated according to the previous annotation of the transcriptome and genome-wide mapping of transcriptional start sites (46). **B**. Alignment of NblD homologs from cyanobacteria belonging to four morphological subsections (51); conserved cysteine residues are marked by an arrow. **C**. Western blotting using anti-FLAG antiserum against tagged NblD and reducing and nonreducing conditions for SDS-PAGE. The expression of *nblD* was induced by nitrogen removal (native P_nblD_ promoter) or the addition of 2 µM Cu^2+^ ions (P_petE_ promoter). All samples were loaded in biological duplicates. **D**. Predicted structure model for NblD generated by SWISS-MODEL (59) as a homodimer using the crystal structure of the *E. coli* Cas2 CRISPR protein 4mak.1 as template. The structure is modelled from position 14A to C48, the position of the latter in the two molecules is given for orientation. Lower part: Alignment of NblD (Target) to template, the sequence is rainbow colored from blue (N terminus) to red (C terminus). The symbols indicate predicted secondary structure (G = 3-turn helix, H = α helix, T = hydrogen bonded turn, B = residue in isolated β-bridge, S = bend, C = coil).

The lengths of the 176 homologs identified in this work (**Supplemental Dataset 1**) vary between 41 aa in *Crocosphaera watsonii* WH 0005 and 121 aa in *Phormidesmis priestleyi* Ana. The alignment of selected NblD homologs highlights the presence of four conserved cysteine residues (**Fig. 1B**). Moreover, these four cysteine residues are conserved also in all other detected homologs except in 7 very short forms, which lack the first cysteine pair (**Supplemental Dataset 1**). The first pair of cysteines is arranged in a CPxCG-like motif, typical for zinc-finger structures in small proteins of bacteria and archaea (57, 58). These structures can bind metal ions and initiate loop formation, which is relevant for transcription factors. Additionally, protein-protein interaction conditioned by sulfur bonds between two cysteines is possible. To test this, we used two strains overexpressing NblD fused to a C-terminal 3xFLAG tag, one under the control of its native P_*nblD*_ promoter and the other under the control of the copper-inducible P_*petE*_ promoter on plasmid pVZ322, yielding strains P_*nblD*__*nblD*3xFLAG and P_*petE*__*nblD*3xFLAG (49). When analyzing total protein extracts from these strains by western blot analyses, we noticed a second band of higher molecular mass under nonreducing conditions but not in the presence of DTT and β-mercaptoethanol (**Fig. 1C**). This result supported interaction, either as a homodimer or with another partner. Consisting with this result, the prediction tool SWISS MODEL (59) modelled NblD as a homodimer and predicted a helical segment in the most conserved part of the protein (**Fig. 1D**).

We conclude that *nblD* genes exist in a wide range of cyanobacteria and in chromatophore genomes of photosynthetic *Paulinella* and that the NblD protein can form dimers, might interact with other biomolecules and that a regulatory role could not be excluded.

### Transcriptomic analysis identifies a functional response to nitrogen step-down in the Δ*nblD* mutant

To identify possible effects on the regulation of gene expression, the *nblD* gene was replaced by a kanamycin resistance cassette and selected to homogeneity (**Fig. S1**). Total RNA was isolated from Δ*nblD* and the wild type immediately before (0 h) and 3 h after the induction of nitrogen depletion to evaluate possible direct effects of NblD on transcription. For the genome-wide assessment of steady-state RNA levels, high-density microarrays were used that employ probes for all 8,916 previously detected transcripts originating from loci on the chromosome or seven plasmids (46, 48). The array design allows the direct hybridization of total RNA, avoiding the pitfalls of cDNA synthesis. As expected, the lack of *nblD* transcription was readily detected in the microarray, indicated by a log_2_ fold change (FC) of -4.5 in the direct comparison between the expression in the wild type and Δ*nblD* after 3 h of nitrogen deprivation (**Table 2**). We performed northern hybridizations to verify the transcriptomic data. While no signal was detected in Δ*nblD* confirming the completeness of gene deletion, an *nblD* transcript was induced upon nitrogen starvation in the wild type, yielding a single band of ∼500 nt (**Fig. 2A**). This matches the sum of the previously calculated lengths of transcriptional unit (TU) 728 and TU731 together, extending from position 729645 to 730159 on the forward strand (GenBank accession no. NC_000911) (46).

**Table 2.**
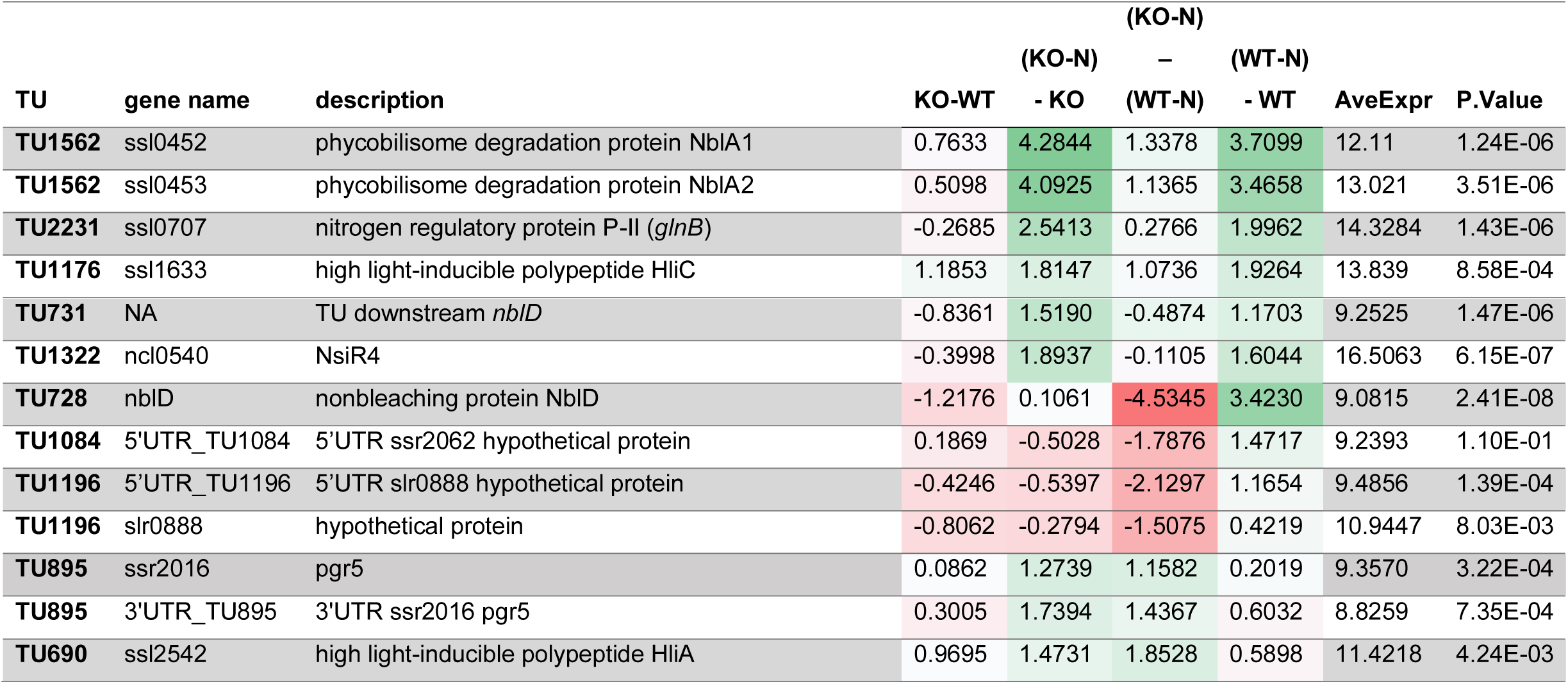
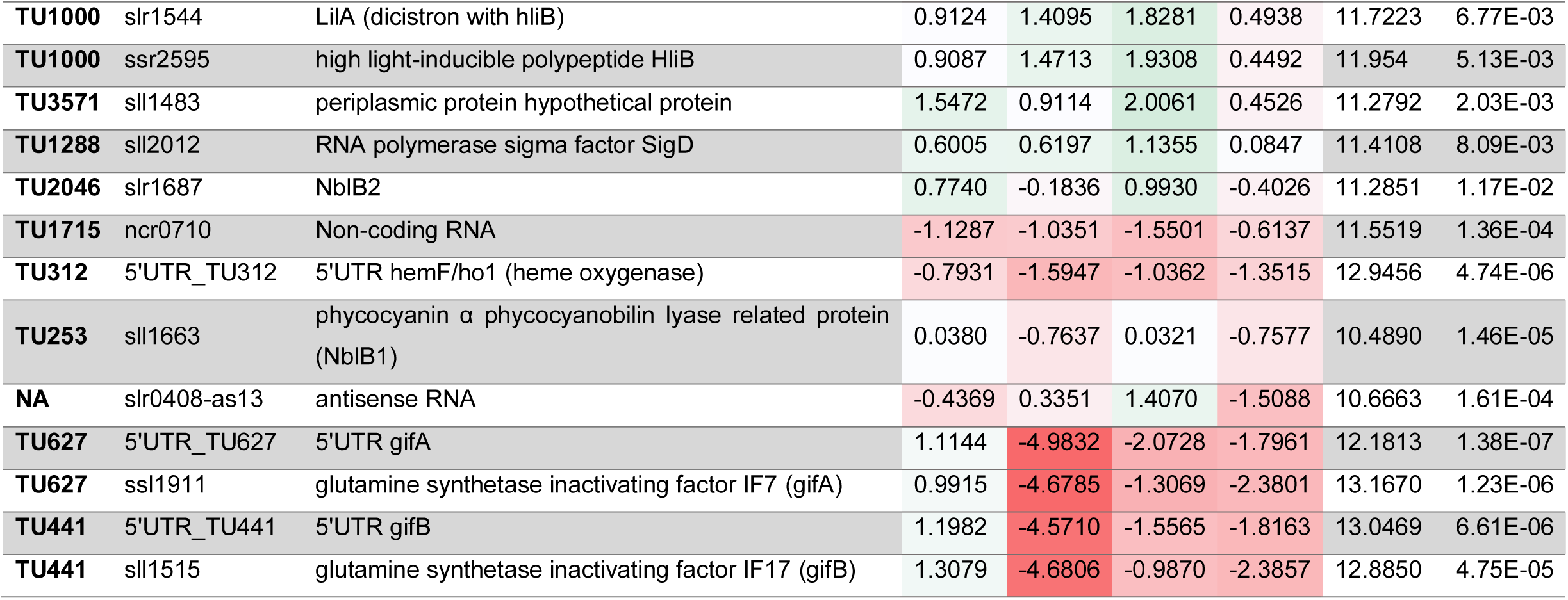
Microarray expression analysis of wild-type *Synechocystis* 6803 and *ΔnblD*. Total RNA isolated before and 3 h after the induction of nitrogen depletion (-N). TUs (transcriptional units) were used as previously described (46). The values show log_2_ fold changes (FC) in gene expression in the indicated comparisons between *ΔnblD* (KO) and the wild type (WT). *P*-values were calculated using Benjamini-Hochberg adjustment; average expression (AveExpr) defines the mean of all quantile-normalized median probe intensities of one probe set.

**Fig. 2.**
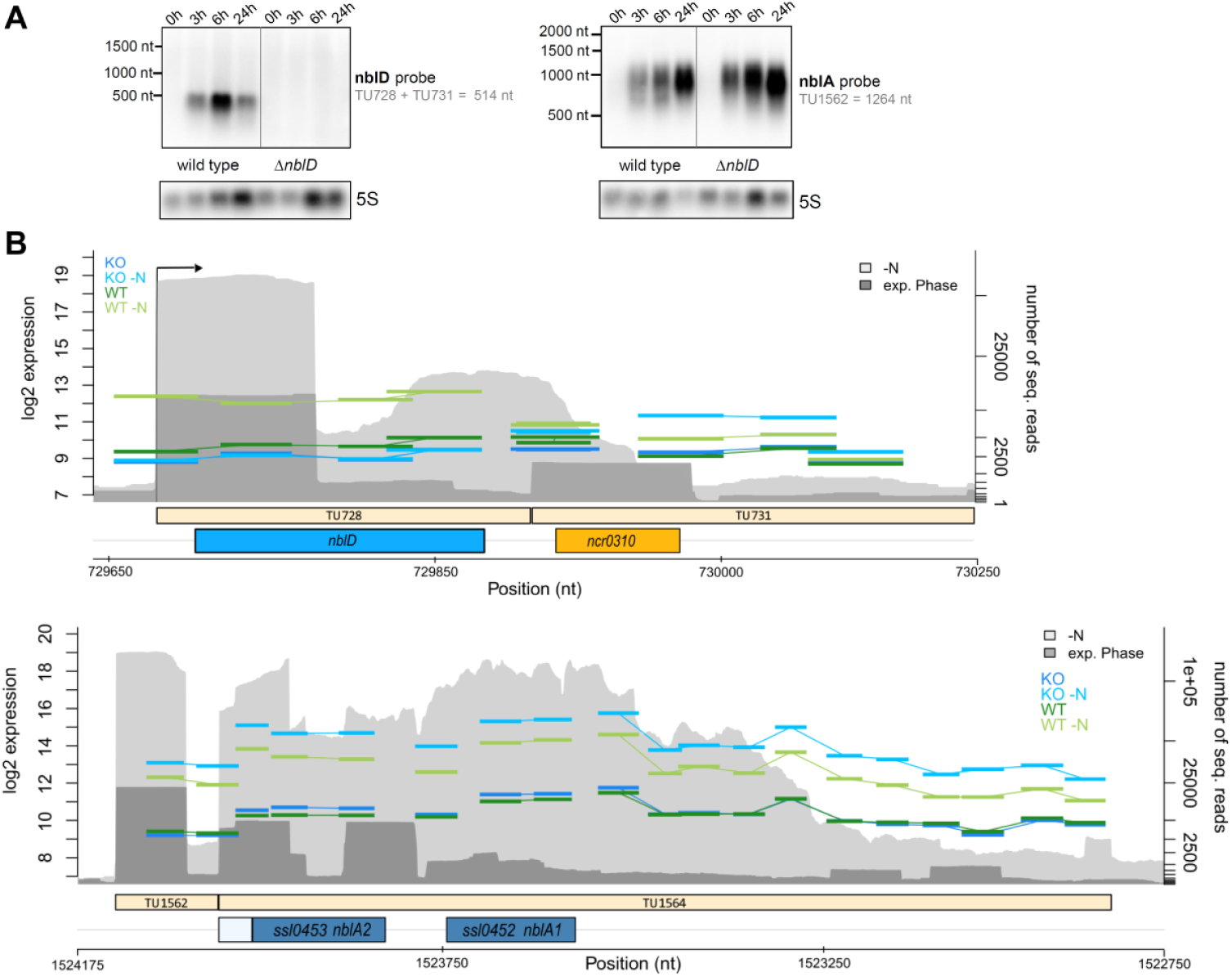
Changes in the abundance of mRNAs in response to an altered nitrogen supply in WT and Δ*nblD*. **A**. Northern blot probing *nblD* and *nblA* in the wild type and Δ*nblD* mutant at 0, 3, 6 and 24 h after the start of nitrogen depletion. **B**. Microarray data visualized for the transcriptional units (TUs) for *nblA* (TU1564) and *nblD* (TU728 + TU731). The log_*2*_ expression values of probes were compared for samples from wild-type (WT) and Δ*nblD* (KO) cultures grown with nitrogen (dark green and dark blue, respectively) and 3 h after the induction of nitrogen depletion (-N, lighter green and lighter blue) at the left scale. The number of sequenced reads (46) for the exponential phase (exp. phase, dark gray) and nitrogen starvation for 12 h (-N, light gray) were included in the background (right scale).

Hence, a transcript with a maximum length of 514 nt contains the 198 nt *nblD* reading frame positioned from nt 729671 to 729871 (including the stop codon). These data yield a 5’UTR of 26 nt and a 3’UTR of 288 nt for *nblD*. The visualization of microarray data at single-probe resolution shows the absence of signals in Δ*nblD* under any condition, a very low basic expression in the wild type under nitrogen-replete conditions that was below the sensitivity threshold of the northern hybridization and an overinduction of the downstream located TU731 in Δ*nblD* under nitrogen starvation (**Fig. 2B**).

Our transcriptomic analysis showed that typical marker genes that become normally induced upon nitrogen step-down were also induced in Δ*nblD* (**Table 2**). For example, the expression of *glnB* encoding the universal nitrogen regulatory protein PII (15, 60) was increased after 3 h of nitrogen starvation in Δ*nblD* by a log_2_FC of 2.5 and in wild type by 2.0 (**Table 2**). NsiR4, a regulatory sRNA that is under NtcA control (61), showed a very similar gene expression change in the mutant and in the control. The most strongly induced genes in the wild type and Δ*nblD* were *nblA1* and *nblA2* with log_2_FCs of 3.5/3.7 and 4.1/4.3, respectively. Their high induction during nitrogen starvation is consistent with previous reports (27). We noticed slightly higher expression of *nblA1A2* in Δ*nblD* than in the control. This effect, as well as the temporally correct and strong induction of transcription in the deletion mutant, was verified by northern analysis (**Fig. 2A**). The detected signal matches approximately the maximum length of the dicistronic *nblA1A2* transcript TU1564 of 1,264 nt (46). Similar to our observation for *nblD*, this transcript is much longer than that needed to encode the small NblA proteins of 60 and 62 amino acids, and this extra length mainly belongs to a very long 3’UTR (**Fig. 2B**).

Some genes, such as *gifA* and *gifB*, both encoding inhibitory factors of glutamine synthetase type I, are strongly repressed upon nitrogen step-down (62). Here, we observed a log_2_FC of -4.7 in Δ*nblD* and -2.4 in the control, both for *gifA* and *gifB* (**Table 2**). Hence, also the reduction in the expression of certain genes in the absence of nitrogen was fully operational in the mutant. Moreover, the amplitudes of expression changes were even stronger in Δ*nblD* than in the wild type, pointing at a possibly more pronounced nitrogen starvation effect in the mutant.

In addition to these marker genes for the short-term response to low nitrogen, we observed a small number of further expression changes, mainly in genes related to pigment biosynthesis or storage, such as *hemF, nblB1, nblB2, hliA, hliB* and *hliC* (**Table 2, Fig. S2**).

Based on transcriptomic analysis, we conclude that the signaling of nitrogen deficiency through the NtcA and PII-PipX systems must have been fully functional in the Δ*nblD* mutant. Hence, the possibility that NblD acted as a regulator or coregulator of their expression could be excluded. **Supplemental Dataset 2** provides a genome-wide graphical overview of probe localization and the corresponding signal intensities for the wild type and Δ*nblD* mutant just before and 3 h following the shift to nitrogen starvation conditions (https://figshare.com/s/308ee7d284599fb2f085). Additionally, the entire dataset can be accessed in the GEO database under the accession number GSE149511.

### Deletion of *nblD* causes a nonbleaching phenotype during nitrogen starvation

To identify a possible phenotype associated with NblD, the Δ*nblD* strain was analyzed. The mutant showed normal growth under nutrient-replete conditions (**Fig. S3**), but the phenotype differed strikingly from the wild type under nitrogen-starvation. The bleaching process was slower and less intense than that of the wild type. Using a complementation plasmid providing an *nblD* gene copy in trans under the control of its native promoter, the wild-type appearance was restored (**Fig. 3A**). Hence, the phenotype of the deletion mutant could be successfully rescued by complementation and the nonbleaching phenotype observed in Δ*nblD* was due to the lack of NblD. Because we could rule out a role of NblD as a transcription factor, we considered its function as a proteolysis adaptor, possibly analogous to NblA. However, ectopic overexpression of *nblD* under the control of the copper-inducible P_*petE*_ promoter did not cause bleaching (**Fig. 3A**), different from the effect of *nblA* overexpression in *S. elongatus* 7942 (63). Therefore, NblD cannot trigger the degradation of the phycobilisome alone, distinguishing it from the activity of NblA, which is a proteolysis adapter protein (31).

**Fig. 3.**
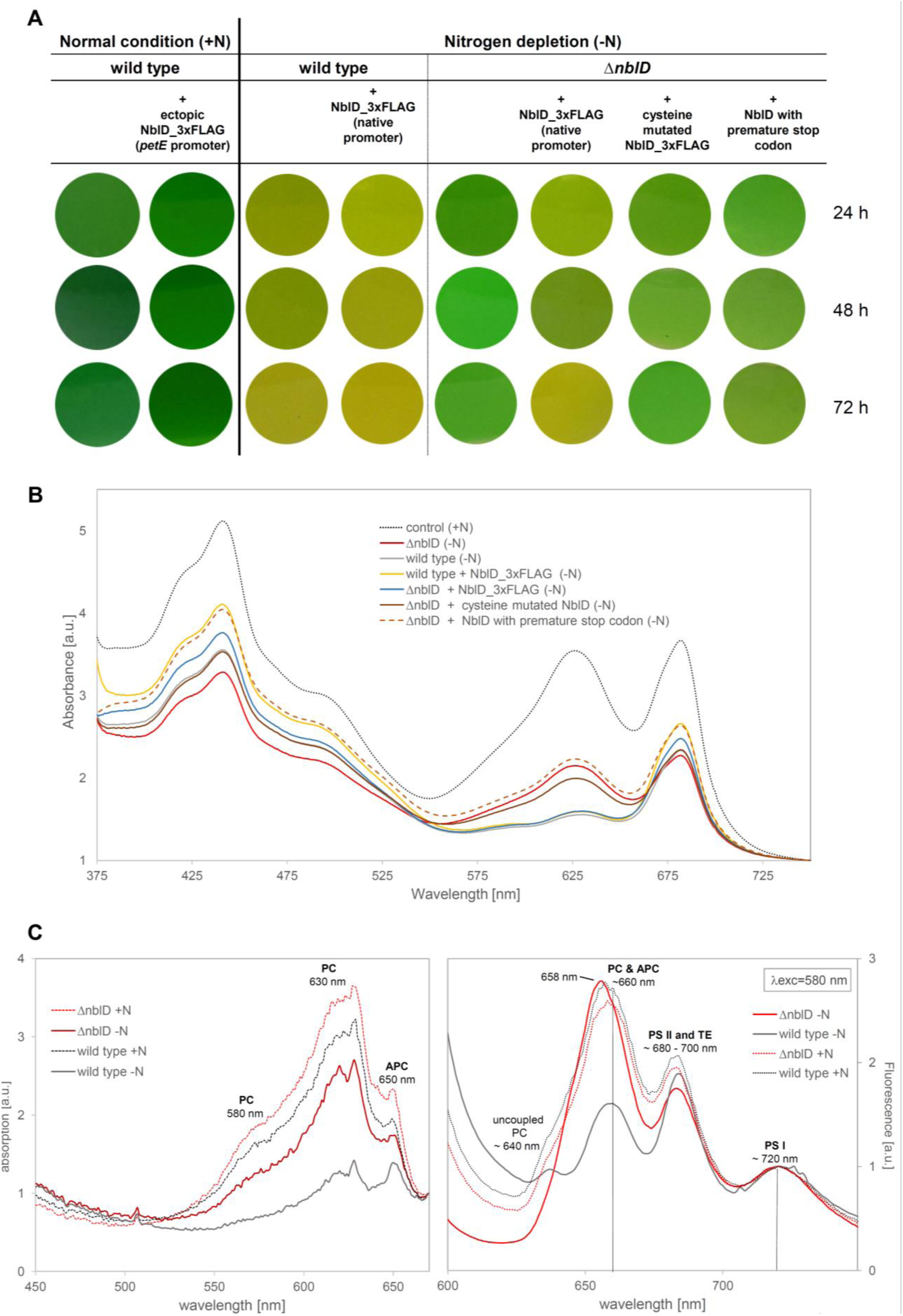
Phenotypical differences between *nblD* mutants and the wild type in nitrogen-replete and -deplete media. **A**. Cultures. **B**. Room temperature absorption spectra for *nblD* mutants normalized to OD_750_. **C**. Low-temperature absorption spectra at 77K for Δ*nblD* and the wild type with and without nitrogen. The spectra were normalized to the minimum at 670 nm. **D**. Emission spectra at 77K and a constant 580 nm excitation for the Δ*nblD* and wild-type strains with and without nitrogen. The spectra were normalized to the photosystem I peak at 720 nm. Acronyms in panels C and D: PC, phycocyanin; APC, allophycocyanin; PSI and PSII, photosystem I and II; TE, terminal emitter.

By replacing the first two cysteines by serine residues (mutations C9S and C12S) we interrupted the first of two cysteine motifs in NblD. Introduction of this construct on a conjugative vector into Δ*nblD* led to only partial complementation, indicating some functional relevance of this first cysteine motif but no absolute requirement for it. Furthermore, an introduced premature stop codon replacing the second amino acid (serine) resulted in a nonbleaching phenotype, indicating that the presence of the translated NblD protein is required for the normal phenotype and not a theoretically possible regulatory feature of the transcript. Moreover, we concluded that the presence of the C-terminal triple FLAG tag did not interfere with the physiological function of NblD because complementation with a FLAG-tagged version of NblD was used to restore the wild-type appearance (**Fig. 3A**).

Consistent with the visual inspection, the different absorption around 630 nm in spectra taken after 48 h of nitrogen starvation indicated that photosynthetic pigments, especially phycocyanin, were still present in the nonbleaching mutants, while these were almost undetectable in wild type and complementation line (**Fig. 3B**). This difference was even more obvious when the data were normalized to the local minimum at 670 nm (**Fig. 3C**). The almost unaffected presence of phycocyanin in Δ*nblD* was confirmed in 77K emission spectra. We used an excitation wavelength of 580 nm, which is the absorption maximum for phycocyanin, and normalized to the photosystem I emission maximum at 720 nm. The emission peak at 660 nm (overlapping peaks of bulk phycocyanin at approximately 652 nm and allophycocyanin at 665 nm) persisted in Δ*nblD* at a high level in the nitrogen starvation condition. Furthermore, this peak was shifted in Δ*nblD* by 2 nm to shorter wavelengths, indicating a different APC-PC ratio in favor of phycocyanin emission. A minor peak at approximately 640 nm indicated the presence of small amounts of remaining uncoupled, monomeric phycocyanin in the wild type. This peak was not detectable in wild type in the nitrogen-replete condition and in Δ*nblD* in neither condition. In comparison, the 680 nm peak, which is caused by fluorescence emitted by photosystem II and the allophycocyanin terminal emitter, was reduced under nitrogen starvation in Δ*nblD* similar to the wild type.

The high amounts of phycocyanin remaining in Δ*nblD* during nitrogen starvation were present in intact phycobilisomes, because these could be isolated 24 h after nitrogen starvation was triggered, whereas wild-type phycobilisomes were already partly dismantled at this time (**Fig. S4**). Taken together, the results indicate a role of NblD in the nitrogen starvation-induced physiological program for the degradation of phycobilisomes during the acclimation to nitrogen starvation.

### NblD interacts with the phycocyanin β subunit

To obtain insight into the molecular mechanism in which NblD is involved, pull-down experiments were performed. The previously constructed *Synechocystis* 6803 P_*nblD*__*nblD*3xFLAG line (49) was used to produce FLAG-tagged NblD fusion protein from a plasmid pVZ322-located gene copy under the control of the native promoter. Three hours after the induction of nitrogen depletion, NblD-FLAG and bound interaction partners were immunoprecipitated from the lysate using anti-FLAG resin and were analyzed by MS (**Table 3**). After the final wash step, the resulting sample was still bound to the M2-anti-flag-resin, showing a dramatic variation in color. While the control samples appeared color-less, the samples containing the NblD-FLAG lysate kept a strong blue color, indicating the presence of likely nondegraded phycocyanin (**Fig. 4A**).

**Table 3.** Most abundant proteins identified by MS analysis for FLAG affinity-pull down samples containing tagged NblD versus knockout samples and their calculated LFQ intensities (84) using MaxQuant. Acronyms: aa, amino acids; MW, molecular weight; PS, photosynthesis; LFQ, label-free quantification. The experiment was performed in biological triplicates, indicated by the numbers 1 to 3.

| Majority protein IDs | GO term | MW [kDa] | aa | ΔnbID 1 | ΔnbID 2 | ΔnbID 3 | nbID_3xFLAG 1 | nbID_3xFLAG 2 | nbID_3xFLAG 3 |
| --- | --- | --- | --- | --- | --- | --- | --- | --- | --- |
| slI1577 phycocyanin $\beta$ subunit | PS antenna | 18.1 | 172 | 4.3E+08 | 3.6E+08 | 2.8E+08 | 1.0E+10 | 1.2E+10 | 7.2E+09 |
| NbID_3xFLAG fusion protein | n/a | 99.3 | 89 | 0 | 0 | 0 | 5.2E+09 | 7.4E+09 | 8.8E+09 |
| slI1578 phycocyanin $\alpha$ subunit | PS antenna | 17.6 | 162 | 5.1E+08 | 2.7E+08 | 2.9E+08 | 4.2E+09 | 5.0E+09 | 5.5E+09 |
| slr0335 phycobilisome core-membrane linker ApcE | PS antenna | 100.3 | 896 | 2.3E+09 | 4.2E+09 | 5.5E+08 | 2.0E+09 | 1.7E+07 | 1.3E+07 |
| slr1140 DegT/DnrJ/EryC1/StrS-family protein | polysaccharide biosynthetic process | 41.6 | 378 | 1.3E+09 | 9.8E+08 | 2.2E+09 | 5.3E+08 | 9.1E+08 | 6.7E+08 |
| slI1580 phycobilisome rod linker polypeptide CpcC1 | PS antenna | 32.5 | 291 | 1.1E+09 | 3.0E+08 | 3.2E+08 | 8.9E+08 | 1.1E+08 | 3.1E+08 |
| slr2067 allophycocyanin $\alpha$ subunit | PS antenna | 17.4 | 161 | 3.4E+08 | 1.4E+08 | 1.6E+08 | 3.5E+08 | 8.9E+07 | 8.2E+07 |
| ssr0482 30S ribosomal protein S16 | ribosome | 95.6 | 82 | 4.0E+07 | 1.7E+08 | 3.0E+08 | 8.7E+07 | 1.1E+08 | 2.7E+08 |
| slI1099 elongation factor Tu | translation | 43.7 | 399 | 4.3E+08 | 6.5E+07 | 1.2E+07 | 2.3E+08 | 4.0E+07 | 1.3E+07 |
| slI1744 50S ribosomal protein L1 | ribosome | 25.9 | 238 | 2.6E+07 | 1.7E+07 | 1.5E+08 | 1.8E+08 | 3.5E+08 | 2.2E+08 |
| slr2051 phycobilisome rod-core linker polypeptide CpcG1 | PS antenna | 28.9 | 249 | 2.4E+08 | 6.5E+07 | 9.3E+07 | 1.7E+08 | 1.1E+07 | 7.5E+07 |
| slI1579 phycobilisome rod linker polypeptide CpcC2 | PS antenna | 30.8 | 273 | 2.6E+08 | 5.7E+07 | 4.6E+07 | 1.3E+08 | 1.6E+07 | 5.6E+07 |
| slr2018 unknown protein | ? | 84.9 | 799 | 1.6E+08 | 8.3E+07 | 2.1E+08 | 1.2E+07 | 1.0E+08 | 0 |
| slr1986 allophycocyanin $\beta$ subunit | PS antenna | 17.2 | 161 | 1.6E+08 | 0 | 3.9E+07 | 1.1E+08 | 2.3E+07 | 9.6E+07 |

**Fig. 4.**
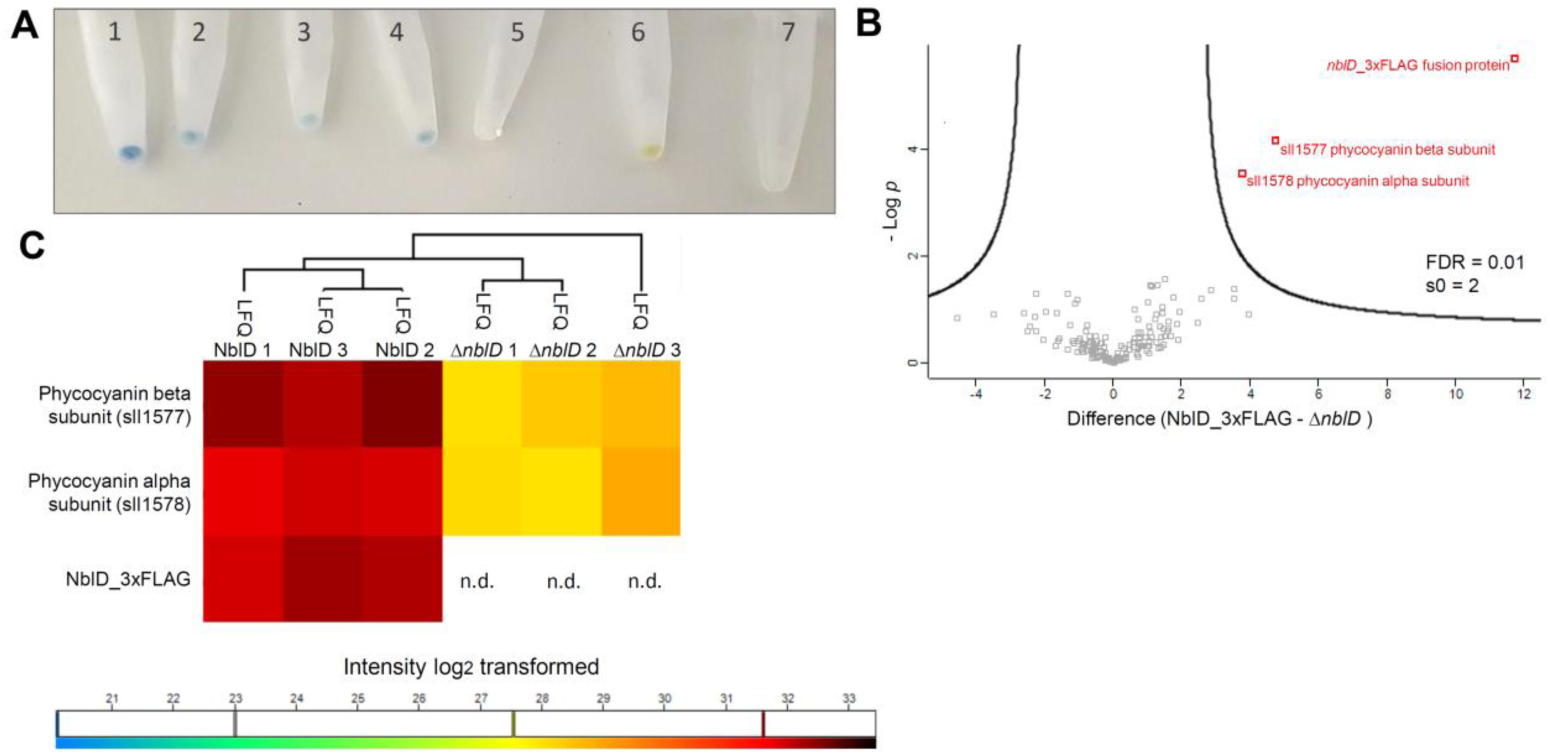
Enrichment of phycobilisome proteins in NblD pull-down experiments. **A**. The collected samples with FLAG-tagged proteins in affinity gels after the final wash step and before the elution were initially incubated as follows: 1–4: lysates containing nblD_3xFLAG; 5: incubated with Δ*nblD* lysate; 6: lysate containing norf1_3xFLAG (49); 7: lysate containing sfGFP_3xFLAG. **B**. Volcano plot of enriched proteins using a false discovery rate (FDR) of 0.01 and s0 (coefficient for variance minimization, see (91)) of 2. **C**. Hierarchical clustering of the most abundant proteins detected in the MS indicating higher log_*2*_ transformed LFQ (label-free quantification) intensities for the phycocyanin subunits in the NblD_3xFLAG containing samples. NblD was not detected (n.d.) in the knockout samples of Δ*nblD*.

Despite its small size, between 7 and 9 of 9 total unique peptides of NblD were detected in these samples by MS analysis. Furthermore, it was highly abundant according to iBAQ values (64), which correspond to the total of all the peptide intensities divided by the number of observable peptides of a protein. The detection of NblD at this early time point, only 3 h after transfer to the low nitrogen condition, was consistent with and extended our observation of high *nblD* mRNA levels at this time (**Fig. 2)**. By contrast, NblD was not detectable in the Δ*nblD* mutant strain. Statistical analysis showed that the phycocyanin α-and β-subunits were the only specifically enriched proteins that coimmunoprecipitated with the NblD-FLAG protein (**Fig. 4B and C**). Comparable results were obtained when the experiment was independently repeated and analyzed at another proteomics facility and after 24 h of nitrogen depletion (data not shown).

To verify the interaction proposed by the results of the MS analysis, we used recombinant NblD fused to a SUMO-6xHis tag at its N terminus in a far western blot approach (65). In this assay, the attached 6xHis tag was then recognized by an HRP-coupled-anti-penta-His-conjugate, providing a signal for proteins bound to the fusion protein (**Fig. 5A**). The signal in the blot overlapped with the phycocyanin β-subunit band at ∼18 kDa, one of the most prominent proteins visible by SDS-PAGE (**Fig. 5B and C**). We included controls to eliminate nonspecific cross reaction of the fusion protein: a *Synechocystis* 6803 mutant lacking all phycocyanins (Δ*cpc*,(66)) and an *E. coli* lysate. In both cases, we did not detect any signal, underlining the specificity of NblD binding to the phycocyanin β-subunit. We conclude from these experiments that NblD specifically interacts with the phycocyanin β-subunit; however, in the coimmunoprecipitation, both subunits were enriched due to the strong natural heterodimer formation between them.

**Fig. 5.**
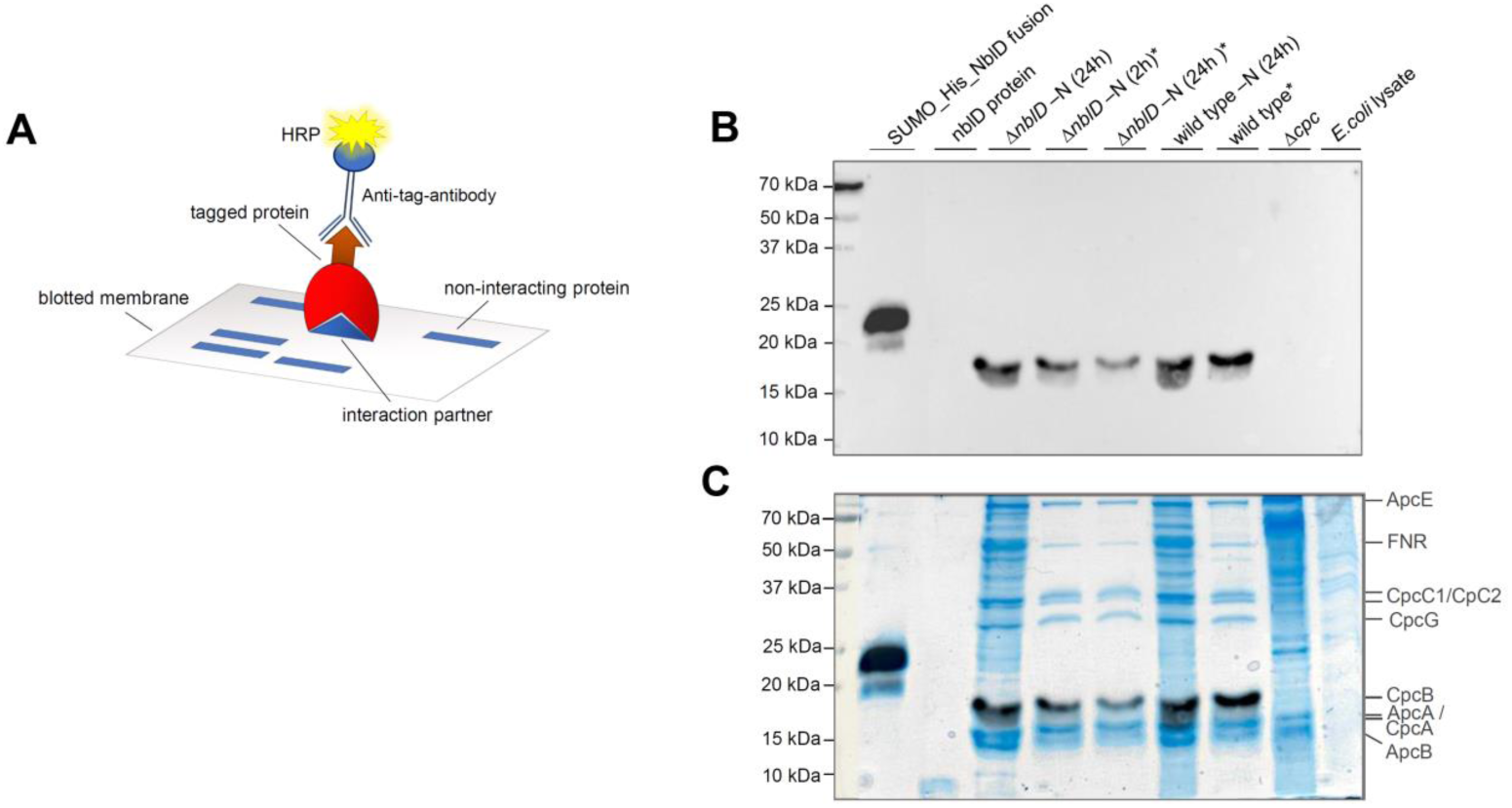
Far western blot analysis. **A**. General principle for the detection of protein-protein interactions by far western blotting using a tagged protein to probe potential interaction partners blotted on a membrane. A secondary antibody, coupled to horseradish-peroxidase (HRP), targets the protein tag. **B**. Western blot signal. **C**. Signal merged to the stained gel to visualize that the signal overlaps with the stained phycocyanin beta subunit. Samples marked with asterisk (*) are isolated phycobilisome proteins.

### The presence of *nblD* is positively selected in competition experiments

We speculated that the lack of NblD might also have a growth effect. However, in short-term growth experiments, strains without the *nblD* gene showed no noticeable growth effect (**Fig. S3**) despite the strong nonbleaching phenotype under nitrogen-starvation conditions (**Fig. 3**). Therefore, we set up a growth competition experiment for a longer period in medium containing only 1 mM NaNO_3_ as the sole source of nitrogen. In this setting, the Δ*nblD* mutant was out-competed by the wild type after 12 generations (**Fig. S5**). These findings directly support the physiological importance of NblD, while its evolutionary conservation suggests that the presence of NblD has been under positive selection in cyanobacteria.

## Discussion

The acclimation of cyanobacteria to nitrogen starvation is a complex physiological process governed by a particular genetic program (10) and involves many proteins and regulatory RNAs with different roles. The main goal of this finely tuned process is a reversible dormant state that is entered until a new source of nitrogen appears. An early major physiological effect is that photosynthesis becomes reduced and, therefore, antenna complexes are diminished. NblD, a small protein is strongly upregulated under nitrogen starvation in *Synechocystis* 6803 (49) and, as shown in this study, participates in phycobilisome disassembly. Other Nbl proteins involved in the disintegration of phycobilisome antenna complexes have been characterized in different cyanobacteria in detail, but most insight has been obtained in *S. elongatus*. The expression of *nblA* genes is strongly induced in both species under nitrogen starvation. NblA was found to interact with the phycocyanin rod antenna structure at a specific groove in the phycocyanin β-subunit (30) but also targeting the allophycocyanin core (67) while NblB was characterized as a chromophore-detaching protein affecting phycocyanin and allophycocyanin (68). The likely ortholog of the *S. elongatus* NblB in *Synechocystis* 6803 is NblB1, whose expression under nitrogen starvation was decreased (**Table 2**), like the *S. elongatus* gene (68). It is noteworthy that knockout mutants of homologs to five *S. elongatus nbl* genes were tested in *Synechocystis* 6803, but only Δ*nblA1* and Δ*nblA2* displayed the non-bleaching phenomenon (32). Later on, Δs*ll1961* was discovered to also yield this phenotype (69). Hence, Δ*nblD* is only the fourth mutant with the *nbl* phenotype in *Synechocystis* 6803. In our coimmunoprecipitation and far western blot analyses, we observed the interaction of NblD with the phycocyanin β-subunit, which is a striking parallel to NblA, the main factor for phycobilisome knockdown by recruitment of a Clp-like protease (31, 70). However, we can rule out a function of NblD as protease adaptor because no protease subunits were found in our interaction screens and its ectopic overexpression under nutrient-replete conditions did not trigger bleaching. We also did not find an interaction with NblA or a likely direct regulatory effect excluding roles as co-effector of NblA or as transcription factor. The only detected interaction was with the CpcB subunit and likely in its chromophorylated state. Phycocyanobilin chromophores are covalently bound to phycocyanin at αCys84, *β*Cys84 and *β*Cys155 and are generally still visible by zinc staining even during SDS-PAGE and in protein pull-down experiments (**Fig. 4A**). These features enabled the far western blot signal for tagged NblD as bait protein bound to the chromophorylated target on the membrane. The chromophores likely also stayed covalently bound to their binding partner in MS enabling the interaction with NblD. Moreover, we found that in the absence of NblD the chromophorylated target remained highly abundant and under nitrogen depletion. These findings point at a pivotal role of NblD in dealing with the phycocyanin linear tetrapyrrole moieties during the early stages of nitrogen starvation. Their uncontrolled release potentially triggers the formation of oxygen radicals and toxic side effects causing redox stress. Indeed, we detected some evidence for this hypothesis in our transcriptome analysis, three of four *hli* genes and the *hliB/scpD* cotranscribed *lilA* (*slr1544*) were induced in Δ*nblD* (**Table 2, Fig. S2**). These genes encode small proteins with an anticipated function in transiently storing chlorophyll molecules during situations causing stress for the photosynthetic apparatus (71–73). Furthermore, the levels of other transcripts encoding proteins known to participate in redox stress responses, like *pgr5* (*ssr2016*) and *sigD* (*sll2012*) were slightly increased in the mutant (**Table 2, Fig. S2**).

According to our data, NblD is a novel factor in the genetic program governing the acclimation response to nitrogen starvation (**Fig. 6**). We propose a hierarchically organized process of phycobilisome degradation, with NblD starting above NblA in *Synechocystis* 6803. In this case, NblD might improve the accessibility to the phycocyanin for NblA. Consequently, if any component is missing in the chain of the nitrogen starvation-triggered disassembly process, a nonbleaching phenotype is obtained. Hence, NblD could interact with the chromophorylated phycocyanin in *Synechocystis* 6803 and bind the phycocyanobilin pigments by disulfide bonds because NblD contains four cysteines arranged in two clusters. We show that a change in the first pair of cysteines results in an appearance similar to the knock-out (**Fig. 3A, B**); thus, an important function of these conserved amino acids can be inferred.

**Fig. 6.**
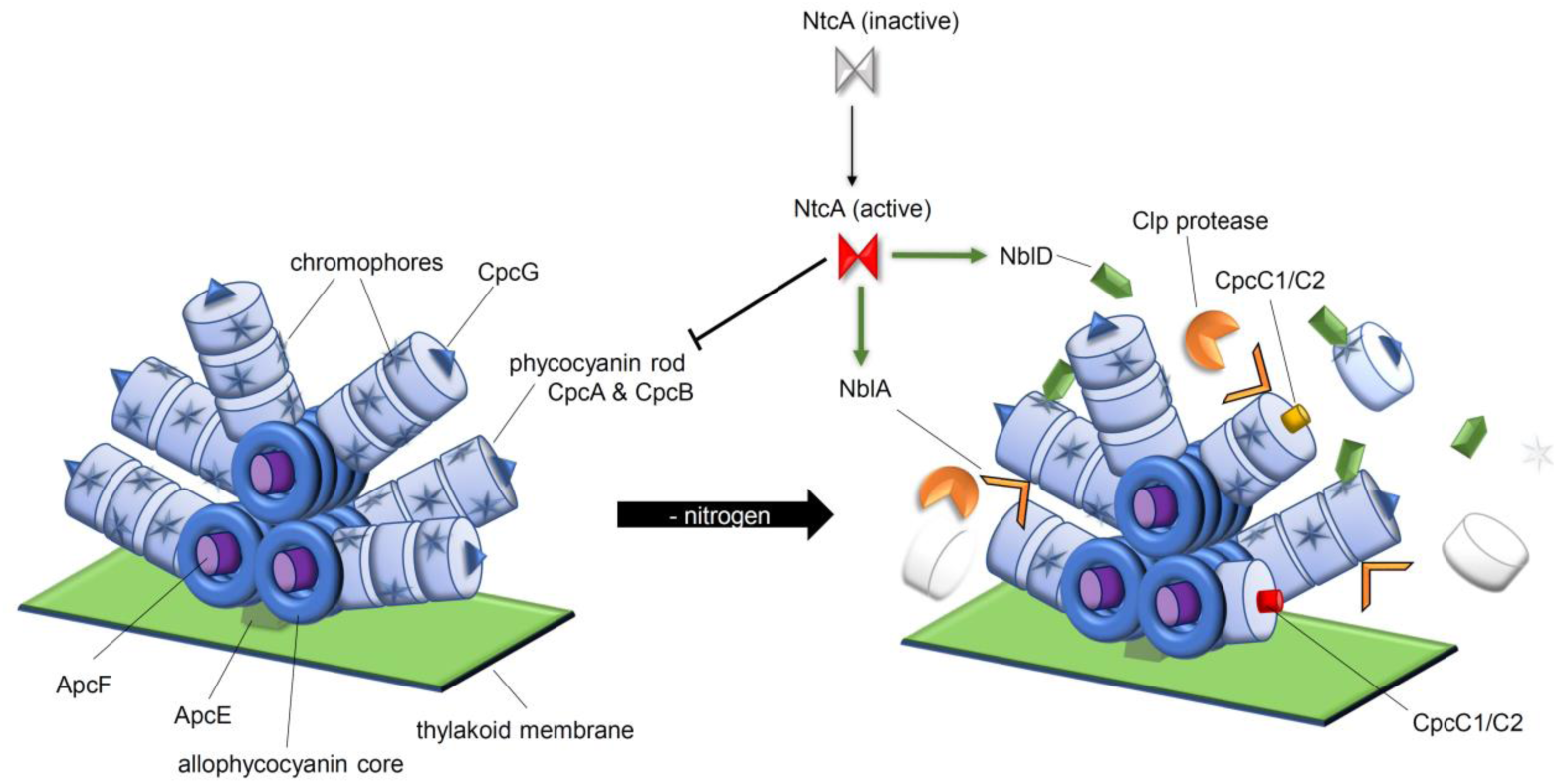
Proposed model for the integration of NblD into the genetic program to respond to nitrogen starvation and its role in phycobilisome degradation in the early phase of nitrogen depletion. NtcA, the major transcriptional regulator of the response to low nitrogen activates transcription of the *nblD* and *nblA* genes (green arrows) while it represses the transcription of the *cpcBAC2C1D* operon (17) encoding the phycocyanin rod and linker proteins. While NblA targets CpcB as protease adaptor recruiting Clp protease, our data suggest that NblD interacts with the chromophorylated phycocyanin and assists the removal of tetrapyrrole chromophores.

We show that NblD is a novel factor in the process that leads to the coordinated dismantling of phycobilisomes. Similar to the NblA proteins that label phycobiliproteins for proteolysis, NblD binds to phycocyanin polypeptides but has a different function. The results show that, even in a well-studied process such as the bleaching response, small proteins can perform crucial functions that have been overlooked thus far.

## Materials and Methods

### Cultivation conditions

Strains were maintained in copper-free BG11 (51) supplemented with 20 mM TES adjusted to a pH of 7.5 at 30°C under continuous white light of 50 µmol photons m^-2^ s^-1^. Mutant strains containing pVZ322 were cultivated in the presence of 50 µg/mL kanamycin and 25 µg/mL gentamycin, while 50 µg/mL of kanamycin was added to cultures of the knockout strain. To deplete cells from nitrogen, the cultures were centrifuged, the cell pellets were washed and resuspended in BG11 without NaNO_3_ (BG11-N). For phenotypic assays, the experimental cultures were grown in medium supplemented with copper lacking antibiotics to prevent any possible effect.

### Construction of mutant and overexpression lines

Using the *Synechocystis* 6803 PCC-M strain (74) as the wild-type and background strain, different mutants were constructed. Strains in which *nblD* expression can be induced via the Cu^2+^-inducible *petE* promoter or controlled via its native promoter on pVZ322-based plasmids have been described previously (49). Knockout mutants were generated by homologous replacement of the *nblD* coding sequence with a kanamycin resistance cassette (*nptII*) and using pUC19 as a vector for subcloning. The construct for gene replacement by homologous recombination was generated by PCR-based AQUA cloning (75) using primers as given in **Table S1**. Total segregation was checked by colony PCR using the primers segregation_nblD_KO fwd/rev and nblD_Km_seq fwd/rev.

To complement the knockout, the self-replicating pVZ322 plasmid encoding different versions of *nblD* were used (primers and vectors are listed in **Tables S1** and **S2**):

1. Encoding *nblD* under control of its native promotor (49) to restore wild type;
2. Same plasmid with a cysteine mutated version of NblD (C9S & C12S);
3. Same plasmid with a premature stop codon (Ser2”STOP”).

The inserts for pVZ322 were assembled into pUC19 by AQUA cloning and then digested, as well as the target pVZ322 vector backbone, for 3.5 h at 37°C by *Xba*I and *Pst*I, producing compatible ligation sites. Fragments were combined and ligated with T4 DNA Ligase (Thermo Scientific) for 4 h at room temperature (RT) and were propagated in *E. coli*. The completed plasmids were introduced into *Synechocystis* 6803 by conjugation (76). The expression levels of *nblD* in the different lines was checked by northern hybridization using a ^32^P-labeled, single-stranded RNA probe produced using primers T7_nsiR6_probe fwd/rev.

### RNA preparation, microarray analysis and northern blot verification

The cultures were starved for nitrogen (-N) for 3 h, 6 h and 24 h to induce *nblD* expression. The time immediately before nitrogen removal (0 h) served as the negative control. After harvesting the culture by filtering through hydrophilic polyethersulfone filters (Pall Supor®-800, 0.8 μm) and immersion in PGTX (77), the samples were snap frozen in liquid nitrogen. RNA isolation was performed as stated previously (78). For northern blot verification, 3 µg of RNA was loaded per well on a denaturing agarose gel and was blotted via capillary blot to Hybond(tm)-N+ membrane (GE Healthcare). The membranes were probed with [α^32^P]UTP-labeled transcripts generated using the MAXIscript® T7 *In Vitro* Transcription Kit (Ambion) and primers T7_nsiR6_probe fwd/rev and T7_nblA_probe fwd/rev as described previously (79). The resulting signal was evaluated by phosphorimaging using a Typhoon(tm) FLA 9500 scanner (GE Healthcare). For microarray analysis, we followed the previously published protocol including direct RNA labeling (80) but using 5 µg of total RNA from the time points 0 h and 3 h -N for wild type and Δ*nblD*. For hybridization, 500 ng Cy3-labeled RNA was applied. The microarray included a duplicate for each sample.

### Protein preparation, proteomic sample preparation and analyses by MS

Cells to prepare total protein samples were collected by centrifugation (3,200 x *g*, 10 min, RT), washed in saline (PBS) supplemented with Protease Inhibitor (cOmplete, Roche) and resuspended in the same buffer. For cell lysis, mechanical disruption using a prechilled Precellys homogenizer (Bertin Technologies) was used. To remove cell debris and glass beads, the culture was centrifuged (1000 × *g*, 5 min, 4°C), and the supernatant was collected for further analysis. Western blots targeting FLAG-tagged proteins were performed using FLAG® M2 monoclonal antibody (Sigma) as described previously (49).

To prepare FLAG-tagged NblD and interacting proteins from total cell lysates and to process mock samples, ANTI-FLAG M2 affinity agarose gels (Sigma) were used. The expression of *nblD* was induced in exponentially growing cultures (800 mL at OD 0.8) by removing nitrogen. After another 3 h of cultivation, the cells were harvested by centrifugation (4000 × *g*, 4°C, 10 min). Cell lysates were obtained as described above (except using FLAG buffer instead of PBS) and then were incubated for 45 min in the presence of 2% n-dodecyl β-D-maltoside to solubilize membrane proteins in the dark at 4°C. After loading the lysate into the packed volume of 100 µL of FLAG agarose on a gravity column (Bio-Rad) and reloading the flow through twice, bound proteins were washed 3 times with FLAG buffer (50 mM HEPES-NaOH pH 7, 5 mM MgCl_2_, 25 mM CaCl_2_, 150 mM NaCl, 10% glycerol, 0.1% Tween-20) and twice with FLAG buffer lacking glycerol and Tween-20.

To achieve maximum reproducibility, protein analyses by MS were performed in two different laboratories and repeated several times. To obtain MS-data, elution was performed using 0.2% RapiGest (Waters) in 0.1 M HEPES pH 8 (MS-grade) and heating for 10 min to 95°C. The RapiGest concentration was decreased to 0.1% by adding 0.1 M HEPES pH 8. The proteins were reduced by incubating in 5 mM dithiothreitol (DTT) and alkylated using 15 mM iodacetamide (IAM) in the dark, each step performed for 20 min at 37°C. Tryptic digestion was performed in two steps; first with 1 µg of trypsin for 2 h at 50°C and second with another 1 µg overnight at 37°C, both shaking at 600 rpm. The peptides were desalted by acidification of the sample to 0.3% TFA final concentration and applying HyperSep C18 tips (ThermoScientific). Thereafter, the peptide concentration was measured using the BCA assay (ThermoScientific). For MS analysis, 500 ng of peptide per sample was analyzed using the EASY-nLC(tm) 1000 UHPLC system (ThermoScientific) coupled to a Q-Exactive plus(tm) Hybrid Quadrupole-Orbitrap(tm) Mass Spectrometer (ThermoScientific) as previously described (81). Raw data were processed and analyzed with MaxQuant (Version 1.6.0.16) using cyanobase (82) data for *Synechocystis* 6803 (Version 2018/08/01) including the small proteins described in reference (49). The proteome raw data acquired by MS were deposited at the ProteomeXchange Consortium (http://proteomecentral.proteomexchange.org) via the PRIDE partner repository (83) under the identifier PXD019019. The intensities were compared using LFQ (label-free quantification) values (84) as illustrated in **Fig. S6** with Perseus (version 1.6.1.3 (85)). In summary, contaminants, reverse sequences and proteins only identified by site were removed from the matrix and LFQ intensities were log_2_-transformed. Before t-test and visualization using a volcano plot, the missing values were replaced by imputation with the normal distribution for each column separately (default settings). For hierarchical clustering (default parameters), only proteins with three valid values in at least one declared group (NblD_3xFLAG and Δ*nblD*) were considered.

### SDS-PAGE and standard western blotting

Proteins were mixed with denaturing and reducing loading dye (5× concentrated: 250 mM Tris-HCl pH 6.8, 25% glycerol, 10% SDS, 500 mM DTT, and 0.05% bromophenol blue G-250). Moreover, 6% ß-mercaptoethanol v/v was added fresh to each sample. The protein samples were separated by either 15% SDS-glycine-PAGE or SDS-tricine-PAGE including the Precision Plus Protein Dual Xtra molecular weight marker (Bio-Rad). To run samples under nonreducing conditions, a native loading dye (4× concentrated: 30% glycerol, 0.05% bromophenol blue G-250, 150 mM Tris-HCl pH 7.5) was used. Gels were stained using either 20 mM zinc sulfate-7-hydrate for reversible zinc staining of chromophores and/or InstantBlue(tm) Coomassie staining (Expedeon). Otherwise, western blots followed a standard procedure as described elsewhere (49).

### Far western blotting

Far western blotting was performed as described previously (65) using purified NblD-SUMO-His-tag fusion protein as the primary antibody. The protein was recombinantly expressed in the *E. coli* ROSETTA strain (Merck) using pE-SUMO as an expression plasmid (86) and was isolated using a 1-mL HisTrap column and Äkta start (GE Healthcare). Before using the fusion protein for far western blotting, it was desalted and concentrated (Vivaspin 20, 10,000 Da MWCO). In the denaturing/renaturing steps of the blotted membrane, milk powder was omitted compared with the protocol provided by Wu *et al*. (65). Between steps, the membrane was washed with TBS-T (20 mM Tris pH 7.6, 150 mM NaCl, 0.1% (v/v) Tween-20). After blocking the renatured membrane with 5% milk powder in TBS-T, 3 µg/mL of fusion protein was used as the ‘primary antibody’ and was incubated at 4°C for at least 6 h. His-Penta-Conjugate (Qiagen) was then used as the secondary antibody (1:5000), targeting the 6×His-Tag of the NblD-fusion protein, with shaking at 4°C for a minimum of 6 h. The membranes were washed in between the single steps at least twice with TBS-T for 5 min at RT. The signals were visualized by applying ECL-spray (Advansta) to the membrane and using a Fusion FX (Vilber) imager.

### Spectrometric measurements

Whole-cell absorption spectra were measured using a Specord® 250 Plus (Analytik Jena) spectrophotometer at room temperature and were normalized to 750 nm. Cultures at an OD_750_ > 1 were diluted with 1× BG11 prior to taking the absorption spectra.

Emission and absorption spectra at 77K were recorded using a FluoroMax (HORIBA Jobin Yvon) spectrofluorometer. If necessary, the cultures were diluted in PBS buffer to avoid saturation effects during measurement. Emission spectra were excited with 580 nm and measured 3 times, and curves were averaged; additionally, absorption spectra were measured 5 times and curves were averaged. In both cases, the slits were set to 5 nm and the integration time was 1 s.

### Phycobilisome isolation

Phycobilisome isolation was performed by adapting a previously described procedure (87). Cells were harvested at exponential OD by centrifugation and were washed once in 0.75 M potassium phosphate buffer (K_2_HPO_4_/KH_2_PO_4_) pH 7 (KP-buffer). Resuspended in the same buffer, the cells were disrupted by two passages through a French pressure cell. Thereafter, the lysate was solubilized with 2% Triton X-100 for 10 min at room temperature with shaking. Debris and insoluble components were removed by centrifugation at 21000 × g for 15 min. The supernatant was loaded onto a sucrose step gradient with 1.5 M, 1 M, 0.75 M and 0.5 M sucrose dissolved in KP-buffer. The blue fraction was collected and precipitated using DOC/TCA precipitation before SDS-PAGE.

### Competition growth assay

According to Klähn *et al*. (61), wild-type and knockout cultures were grown in precultures without antibiotics separately and then were mixed to equal cell numbers in triplicate in 1 mM NaNO_3_ limited BG11 to a final OD_750_ of 0.2. The cell numbers were calculated by microscopic counting and OD_750_ measurement. In parallel, separate control strains of the wild type and Δ*nblD* were grown in limited medium. Every 3 to 4 days, the cultures were diluted to an OD_750_ of 0.2. Additionally, 2.5 µL of 1:10 and 1:100 diluted cultures were spotted on nitrogen-replete BG11 agar plates containing either no antibiotics or 40 µg/mL of kanamycin. The growth of all cultures was documented by scanning the plates using an EPSON scanner. The colonies were counted, and the cell numbers were calculated and compared with controls. Furthermore, the cultures were checked by PCR for their allele composition using the primers *nblD*_Km_seq fwd/rev.

## Acknowledgments

We are very grateful to Tasios Melis, University of California, Berkeley, who kindly provided the phycocyanin mutant for the far western blot analysis. It was a great pleasure to work with your strain! The authors also thank Martin Hagemann (Rostock), Jörg Soppa and Harald Schwalbe (both Frankfurt) and Annegret Wilde (Freiburg) for helpful discussions. Furthermore, we thank Viktoria Reimann for helping with the microarray and Stefan Tuskan for support in setting up the competition assay experiment.

## Author Contributions

WRH designed the project and secured funding. MF, OS, PS and BM carried out MS

-based proteomic analyses. CS supported the microarray visualization and spectroscopic analyses, NFD contributed to the pigment analysis and data interpretation. All other experiments and analyses were performed by VK. VK and WRH wrote the manuscript with input from all authors.

